# Live-imaging of endogenous neurofascins reveals glial adhesion shapes developing nodes of Ranvier

**DOI:** 10.64898/2026.07.24.740328

**Authors:** Phoebe Lyster-Binns, Katy Marshall-Phelps, Sophie Siems, Maira Pyrgioti, Rafael Almeida

## Abstract

The organisation of myelinated axons into specialised domains is essential for saltatory conduction and depends on the polarised distribution of neuronal and glial cell-adhesion molecules, including neurofascins. However, how these domains assemble and refine in vivo remains unclear, as longitudinal imaging of endogenous proteins within intact nervous systems remains challenging. Here, we generated knock-in zebrafish in which endogenous neuronal and glial neurofascins are fused to fluorescent proteins, which enabled live-imaging without perturbing node assembly or introducing overexpression artefacts. Using these reporters, we visualised neuronal and glial neurofascin dynamics, and the formation of nodes of Ranvier and paranodes in vivo. We find that nascent nodes progressively compact in developing peripheral nerves, and that this process depends on glial neurofascin. Our novel toolkit for imaging endogenous neurofascins reveals a glia-dependent mechanism that shapes the developmental refinement of nodal architecture after their initial assembly. Our findings imply that glia-driven modulation of paranodes can be a mechanism for nervous systems to regulate circuit function.

## Introduction

The rapid transmission of information in vertebrate nervous systems depends on a highly ordered molecular architecture along axons. Distinct axonal domains, such as nodes of Ranvier where sodium channels concentrate and the adjacent paranodes that form tight axo-glial junctions, are essential for saltatory conduction. Their precise geometry and spatial distribution along the length of an axon modulate action potential propagation (Hartline and Colman, 2007). Recent studies highlight not only the diversity of axonal domain distributions in vertebrate nervous systems, but also their lifelong plasticity and involvement in disease (Almeida and Lyons, 2017; Bacmeister et al., 2020; Bacmeister et al., 2022; Hill et al., 2018; Tomassy et al., 2014). How these domains are established during development and regulated throughout life is therefore a central question in understanding how neurons and glia cooperate to sculpt functional neural circuits.

A family of molecules that act as core organisers of axonal domains are neurofascins, immunoglobulin cell adhesion molecules that orchestrate the assembly and maintenance of nodes and paranodes. In mammals, a single *NFASC* gene encodes neuronal splice isoforms, such as *NF186*, and a glial isoform, *NF155* (Davis et al., 1996; Volkmer et al., 1992). *NF186* localises at nodes and induces sodium channel clustering; while *NF155* concentrates at paranodes and is a key component of the adhesion complex that attaches myelin to the axon, providing a diffusion barrier between domains (Fig. 1A) (Sherman et al., 2005; Zonta et al., 2008). Analysis of endogenous neurofascin proteins has relied mainly on immunohistochemistry, which captures only static distributions of proteins in fixed tissue. Transgenic overexpression of tagged constructs, another approach employed in some in vitro and in vivo studies (Klingseisen et al., 2019; Malavasi et al., 2021; Vagionitis et al., 2022; Zhang et al., 2012) could in principle subtly alter the normal morphology or physiology of these submicron domains (Dumitrescu et al., 2016; Hedstrom et al., 2007). In addition, independently manipulating only neuronal or glial neurofascin requires sophisticated genetic targeting of individual splice isoforms (Pillai et al., 2009; Sherman et al., 2005; Zonta et al., 2008). Therefore, it has been difficult to ascertain the real-time dynamics and plasticity of endogenous neurofascins and of axonal domains during the development of myelinated axons in vivo. What is lacking are faithful reporters for live, real-time, longitudinal, non-perturbative visualisation of endogenous neurofascins and axonal domains at single-axon resolution.

**Figure 1.**
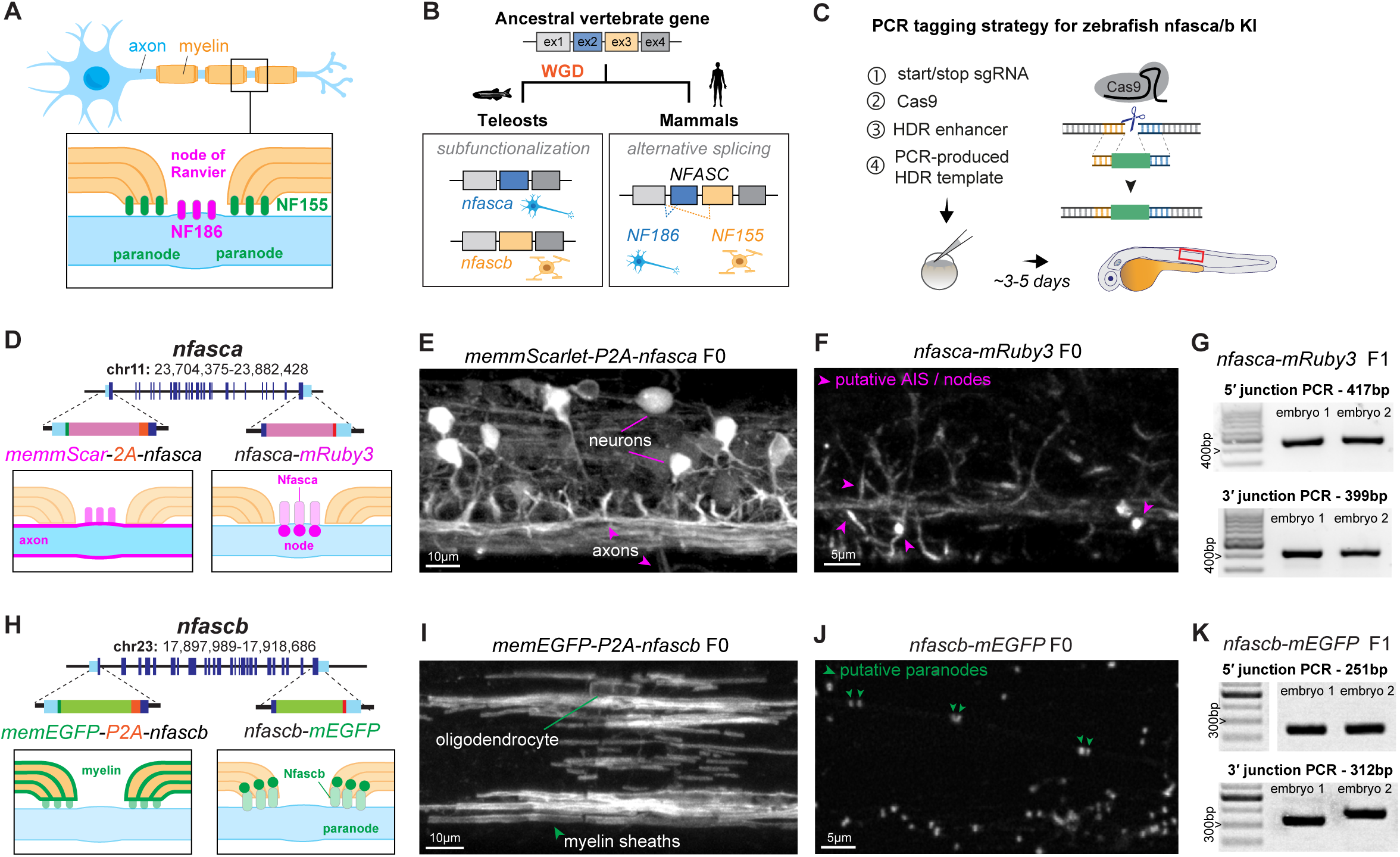
Design and production of novel *nfasca/b* knock-in alleles. (A) Molecular organisation of mammalian neurofascins along axonal domains. (B) Whole Genome Duplication (WGD) in teleosts created separate *NFASC* paralogues in zebrafish. (C,D,H) Design of N- and C-terminal *nfasca/b* knock-ins, with predicted labelling highlighted. Boxed region in C indicates location images in E, FI, J). (E,I) Lateral spinal cord view in 4-5dpf embryos injected with N-terminal, cellular tagging knock-in reagents. (F,J) Lateral spinal cord view in 4-5dpf embryos injected with knock-in reagents for C-terminal protein tagging. (G,K) Junction PCR analysis (and Sanger sequencing, Fig S1) show precise C-terminal integration in F1 embryos.

Zebrafish are a useful vertebrate model for live-imaging the nervous system in a non-invasive manner. Following a whole-genome duplication event, zebrafish possess two separate neurofascin paralogue genes, *nfasca* and *nfascb*, with protein domain organisation similar to *NF186* and *NF155* respectively (Klingseisen et al., 2019; Vagionitis et al., 2022). This divergence provides a unique opportunity to independently analyse neuronal and glial neurofascin in vivo. Here, we describe two novel zebrafish lines, *nfasca-mRuby3* and *nfascb-mEGFP*, generated by precise in-frame knock-in of fluorophore coding sequences at *nfasca/b* genomic loci to tag endogenous neuronal and glial neurofascins respectively. Using these tools, we monitored the developing posterior lateral line nerve, a peripheral mechanosensory nerve in the zebrafish embryo easily accessible for highly resolved live-imaging of myelinated axons and their domain organisation. We show that these live reporters faithfully label nodes and paranodes without disrupting axonal domain organisation, and that they avoid the morphological artifacts produced by overexpression of reporters. Through longitudinal imaging of the same individual nerves, we also identify compaction of nodal domains over time that is dependent on glial neurofascin.

## Results

### Zebrafish neurofascin paralogues show distinct neuronal and glial expression patterns

We first sought to confirm that zebrafish neurofascin paralogues, *nfasca* and *nfascb*, acquired the expression patterns of the mammalian splice isoforms *NF186* and *NF155*. Following whole genome duplication, paralogues can diverge in exon structure and expression pattern to recapitulate alternative splice isoforms of the ancestral gene, an evolutionary process known as subfunctionalization (Fig. 1B) (Lambert et al., 2014).

To confirm *nfasca/b* expression patterns, we first used our PCR-tagging strategy (Zhang et al., 2023) to knock-in membrane-targeted fluorescent tags in *nfasca/b* (Fig. 1C,D,H), enabling us to live-image the morphology of expressing cells to identify them. To label Nfasca^+^ cells, we injected fertilised eggs with a *nfasca* start-codon targeting sgRNA, Cas9 and an HDR template with *nfasca* homology arms flanking a membrane-tethered mScarlet and ribosome-skipping P2A peptide sequence. Correct integration should lead *nfasca*-expressing cells to produce memmScarlet separately from intact endogenous Nfasca. 4/155 (∼3%) embryos screened at 3-4dpf had memmScarlet expression. This localised in cell bodies with a neuronal morphology and position in the zebrafish spinal cord, as well as in their dendrites and axons, and there was no obvious glial expression (Fig. 1E). In parallel, to label Nfascb^+^ cells, we injected wildtype eggs with a *nfascb* start-codon targeting sgRNA, Cas9 and a memEGFP-2A HDR template flanked by *nfascb* homology arms. 5/70 (7%) embryos screened at 5dpf had memEGFP expression. This localised in cells with a clear myelinating oligodendrocyte morphology in the spinal cord, and there was no apparent neuronal expression (Fig. 1I). These knock-in efficiencies are in line with our previous study (Zhang et al., 2023). These data confirm zebrafish *nfasc* orthologs compartmentalised neuronal (*nfasca*) and glial (*nfascb*) expression patterns.

### Generation of knock-in alleles for live-imaging of endogenous Nfasca/b proteins

We then sought to tag endogenous Nfasca/b proteins for direct live-imaging of their localisation in real time in the developing nervous system. Neuronal *NF186* localises to the axon initial segment and nodes of Ranvier; and glial *NF155* localises to the myelin paranodal loops (Fig. 1A). The node and paranode-organising functions of *NF186* and *NF155* are mediated through interactions in their extracellular domains (Chataigner et al., 2022; Eshed et al., 2007). To minimise perturbation of endogenous Nfasca/b functions, we therefore chose to target their intracellular C-termini, introducing a short linker to facilitate correct protein folding. We used our PCR tagging strategy to integrate spectrally distinct, monomeric reporter sequences before the *nfasca/b* stop codons (Fig. 1D-H), to enable simultaneous Nfasca/b imaging: mRuby3 for Nfasca and monomeric EGFP (mEGFP) for Nfascb. To tag the Nfasca C-terminus, we co-injected a *nfasca* stop-codon targeting sgRNA, Cas9, and an HDR template with *nfasca* homology arms flanking the mRuby3 sequence. This led to detectable mRuby3 signal in the nervous system of 12/193 (∼6%) injected embryos, which appeared as linear ∼5-20µm segments, putatively axon initial segments or other unmyelinated regions; and as more punctate clusters, potentially nodes of Ranvier (Fig. 1F, arrowheads). After raising these embryos, we recovered a *nfasca-mRuby3* allele with an efficiency of 33% - 1 of 3 screened adult F0 animals transmitted to 31% of its offspring. PCR with junction-specific primers (Fig. 1G) and Sanger sequencing (Fig. S1) validated on-target, precise integration in our novel *nfasca-mRuby3* allele.

To tag the Nfascb C-terminus, we co-injected a *nfascb* stop-codon targeting sgRNA, Cas9, and an HDR template with *nfascb* homology arms flanking the mEGFP sequence. This led to detectable mEGFP signal in the nervous system of 6/131 (∼5%) injected embryos, which appeared as individual or paired bright clusters, putatively paranodes (Fig. 1J, arrowheads). We recovered *nfascb-mEGFP* alleles with an efficiency of 67% - 2 of 3 screened adult F0 animals transmitted to 6-23% of their offspring. PCR with junction-specific primers (Fig. 1K) and Sanger sequencing validated on-target, precise integration in one of our novel *nfascb-mEGFP* alleles; and showed the other allele had a partial 52bp duplication of the 3’ homology arm after the stop codon, in the 3’ UTR, not predicted to affect protein function (Fig. S1). Homozygous *nfasca-mRuby3* and homozygous *nfascb-mEGFP* knock-in embryos developed normally and were indistinguishable from wildtype animals.

### Expression patterns of endogenous Nfasca/b in myelinated tracts

To directly examine endogenous Nfasca/b proteins in vivo, we live-imaged *nfasca-mRuby3* or *nfascb-mEGFP* knock-in embryos between 2-5dpf, carrying the established *Tg(mbp:EGFP-CAAX)* or *Tg(sox10:mRFP)* transgenes respectively to visualise myelin (Almeida et al., 2011; Kirby et al., 2006). We focused on the hindbrain and spinal cord in the central nervous system (CNS), and on the posterior lateral line nerve (pLL) of the peripheral nervous system (PNS). The pLL is a compact, superficial nerve localised along the length of the zebrafish trunk that is well suited for imaging sparse or individual axons.

Nfasca-mRuby3 signal localised diffusely in neuronal structures at 2dpf, prior to myelination, including in Rohon-Beard soma-like profiles in the dorsal spinal cord, and along axonal tracts (Fig. S2A). At 5dpf, after the onset of both CNS and PNS myelination, Nfasca-mRuby3 signal concentrated at more discrete axonal segments, including axon-initial segment like profiles and nodes of Ranvier flanked by adjacent mbp:EGFP-CAAX^+^ myelin sheaths (Fig. 2A-D). We also observed heminode-like clusters, flanked only on one side by a single myelin sheath (Fig. 2B). At 5dpf, consistent with a *NF186*-like function at nodes of Ranvier, essentially all gaps between adjacent myelin sheaths were Nfasca-mRuby3^+^ in both the CNS (83/84 gaps sampled in 3 animals) and PNS (24/24 gaps sampled in 6 animals). Higher magnification imaging of the pLL nerve in *nfasca-mRuby3* embryos (Fig. 2C), including in embryos carrying the axonal membrane reporter *Tg(nbt:axonGCaMP7s)* (Fig. S2B), showed mRuby3 signal present along non-nodal regions along the axon (see also Supplementary Movie S1). This is consistent with a previously described trafficking population of NF186 molecules (Bekku and Salzer, 2020; Ghosh et al., 2020). Time-lapse imaging combined with photobleaching of nodal areas indeed revealed dynamic movement of non-nodal Nfasca-mRuby3^+^ particles along the axon (Fig. S2C).

**Figure 2.**
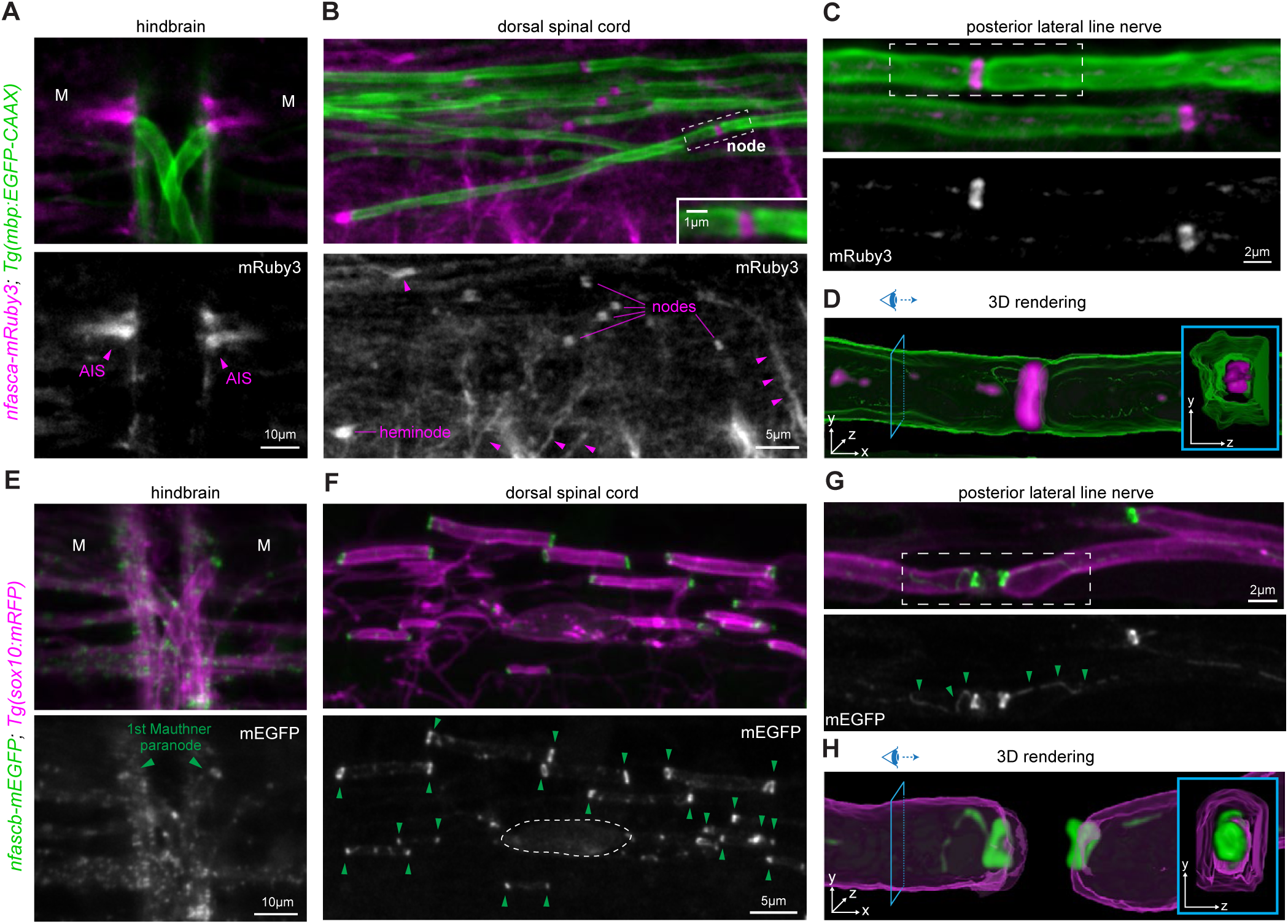
Localisation of endogenous Nfasca/b proteins relative to myelin reporters. (A,E) Dorsal, hindbrain views of 4dpf *nfasca/b* KI crossed with myelinating cell reporters. Mauthner soma location indicated by M. (B,F) Lateral, dorsal spinal cord views of 5dpf *nfasca-mRuby3* and 3dpf *nfascb-mEGFP* crossed with myelinating cell reporters. In B, Nfasca^+^ nodes and heminodes adjacent to myelin sheaths are highlighted, and unmyelinated Nfasca^+^ axonal segments (arrowheads). In F, dashed line indicates oligodendrocyte cell body, and arrowheads highlight Nfascb^+^ edges of all sox10:mRFP^+^ myelin sheaths. (C,G) High-magnification lateral views of pLL in 5dpf *nfasca-mRuby3* and 4dpf *nfascb-mEGFP* animals. Arrowheads in G indicate Nfascb-mEGFP at the inner tongue. (D, H) 3D surface rendering of myelin (green in D, magenta in H) and Nfasca (magenta in D) or Nfascb (green in H) corresponding to boxed regions in C and G. Insets show orthogonal zy views at positions indicated by the eye icon.

Nfascb-mEGFP signal was not detectable at 2dpf, prior to the onset of myelination; while at 4dpf, Nfascb-mEGFP signal concentrated at discrete clusters localised at the ends of sox10:mRFP^+^ myelin sheaths in both CNS and PNS (Fig. 2E-H). At 4-5dpf, consistent with a NF155-like function at paranodal junctions, nearly all myelin sheath ends were Nfascb-mEGFP^+^ in both the CNS (236/249 sheath ends sampled in 6 animals) and PNS (228/228 sheath ends sampled in 8 animals). In addition, higher magnification imaging of the pLL nerve also revealed a fraction of mEGFP signal present along the length of myelin sheaths, consistent with localisation at the inner tongue (Fig. 2G-H, Supplementary Movie S2). Time-lapse imaging of the pLL at 4dpf, when myelination is progressing, also revealed clear dynamics of peripheral Nfascb-mEGFP^+^ clusters, such that even a ‘mature’ configuration of two paranodes flanking a nodal gap can undergo rapid displacement along axons, while others remain stable (Fig. S2D).

Collectively, our findings are consistent with previous studies suggesting Nfasca/b being the zebrafish orthologs of *NF186/NF155,* respectively (Klingseisen et al., 2019; Vagionitis et al., 2022), and establish our precisely tagged *nfasca-mRuby3* and *nfascb-mEGFP* lines as powerful tools for longitudinal live-imaging of endogenous neuronal and glial neurofascins in vivo.

### Fluorescent tags do not perturb nodal Na_v_ channel clustering

To determine whether the presence of the mRuby3 or mEGFP tags perturbs endogenous Nfasca or Nfascb functions, we examined node of Ranvier formation by assessing the expression of voltage-gated sodium channels along pLL axons. In 5dpf *nfasca-mRuby3* or *nfascb-mEGFP* homozygous larvae, immunostaining for voltage-gated sodium channels (NaCh) and acetylated tubulin (AcTub) showed nodes and axons, respectively, similar to wildtype animals. In both genotypes, NaCh immunoreactivity was observed as discrete puncta distributed along the length of labelled axons (Fig. 3A), consistent with the localisation of sodium channels at nodes of Ranvier. NaCh-positive puncta appeared as tightly clustered domains (Fig. 3A), at similar frequency along wildtype and homozygous knock-in nerves (Fig. 3C). We additionally tested if Schwann cells myelinated normally when Nfascb was tagged with mEGFP, by imaging mosaically labelled Schwann cells in the pLL in wildtype and homozygous *nfascb-mEGFP* animals and assessing their length (Fig. 3B). We found no significant difference in mean sheath length between genotypes, assessed at 5dpf (Fig. 3D). Together, the lack of detectable differences in the emergence of NaCh clusters, Schwann cell myelin sheath length, or survival between homozygous knock-in and wildtype animals suggest that endogenous Nfasca and Nfascb functions as organisers of axonal domains are not disrupted by the fluorescent tags.

**Figure 3.**
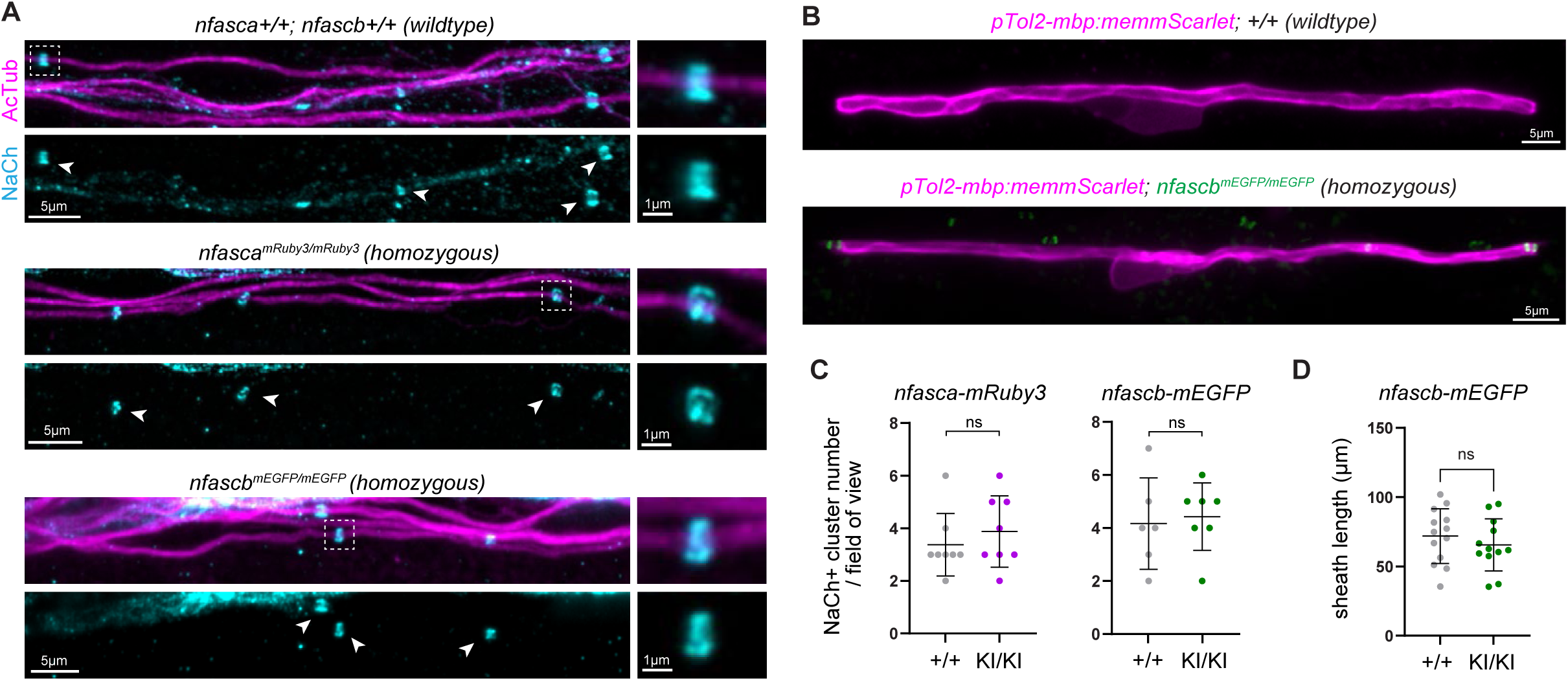
Novel *nfasca/b* knock-in alleles do not disrupt normal node formation. (A) Lateral views of the pLL in wildtype and homozygous *nfasca-mRuby3* and *nfascb-mEGFP* 5dpf animals immunostained for sodium channel (NaCh) and acetylated tubulin (AcTub). Exemplar nodes indicates by arrowheads and dashed heads, magnified in side panels. (B) Individual Schwann cells in the pLL of 5dpf +/+ and *nfascb-mEGFP* KI/KI animals. (C) pLL node density comparison across genotypes: p = 0.446, unpaired t-test, 8 wildtype (+/+) and 8 *nfasca-mRuby3* homozygous (KI/KI) animals; p = 0.759, unpaired t-test, 6+/+ and 7 *nfascb-mEGFP* KI/KI animals. (D) Schwann cell length comparison: p=0.421, unpaired t-test, 13 cells in 13 +/+ and 13 cells in 12 KI/KI animals.

### Knock-ins circumvent conventional reporter overexpression artefacts

We then assessed how our knock-ins compared to conventional transgenic overexpression of tagged neurofascin proteins, a labelling approach which we previously found to perturb the subcellular localisation of tagged proteins in neurons (Zhang et al., 2023). We compared *nfasca-mRuby3* animals with animals injected with a UAS:nfasca-mRuby3 construct expressed in individual pLL neurons, using the *Tg(KalTA4u508)* driver line (Antinucci et al., 2019). At 5dpf, both groups were imaged at multiple points along the pLL, with axons labelled using the *Tg(nbt:axonGCaMP7s)* reporter (Fig. 4A-B). Discrete Nfasca-mRuby3^+^ clusters could be observed along axons in both conditions with a typical nodal morphology. To assess whether node morphology differed between the two approaches, we analysed fluorescence intensity profiles of individual Nfasca-mRuby3^+^ clusters. Average node length was significantly increased in the overexpression condition compared to the knock-in line (Fig. 4E). Additionally, diffuse Nfasca-mRuby3^+^ signal was also observed along axons in overexpressing animals (Fig. 4B, small arrowheads).

**Figure 4.**
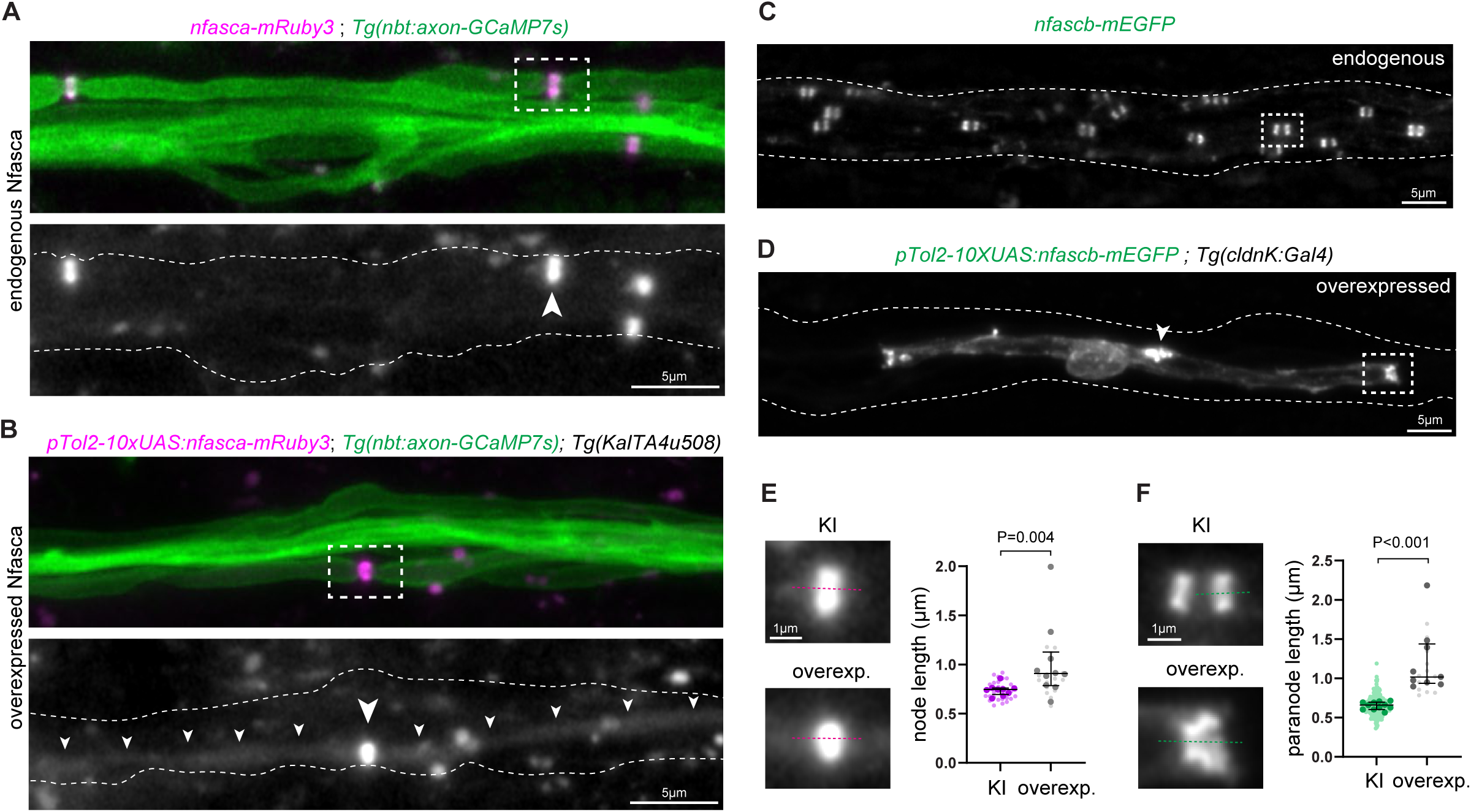
Fluorescently tagging endogenous neurofascins circumvents overexpression artefacts. (A-B) Live-imaging of endogenous Nfasca-mRuby3 and Nfasca-mRuby3 overexpression in the pLL. Nodes indicated by large arrowheads and non-nodal signal in overexpressing cell by small arrowheads. Boxed exemplar nodes magnified in I. (C-D) Live-imaging of endogenous Nfascb-mEGFP and Nfascb-mEGFP overexpression in the pLL. Arrowhead in H indicates non-paranodal signal in overexpressing cell. Boxed exemplar paranodes magnified in J. (E) Node length comparison: p = 0.004, Mann-Whitney test, 9 *nfasca-mRuby3* and 10 *UAS:nfasca-mRuby3* overexpressors. (F) Paranode length comparison: p<0.001, Mann-Whitney test, 11 *nfascb-mEGFP* and 9 *UAS:nfascb-mEGFP* overexpressors.

We also compared the *nfascb-mEGFP* line with animals injected with a UAS:nfascb-mEGFP overexpression construct expressed in myelinating Schwann cells, using a *Tg(cldnK:Gal4)* driver line (Münzel et al., 2012). At 5dpf, both groups were imaged along the pLL, and discrete mEGFP clusters could be observed consistent with paranodal localisation at the edges of myelin sheaths in both conditions (Fig. 4C-D). To assess whether Nfascb cluster morphology differed between the two approaches, we analysed fluorescence intensity profiles of individual Nfascb-mEGFP^+^ clusters. Average paranode length was significantly increased in the overexpression condition compared to the knock-in line (Fig. 4F). Additionally, in 8/9 overexpressing Schwann cells, other focal regions of high mEGFP signal were present near the Schwann cell soma and along the myelin sheath (Fig. 4D, arrowhead). Together, these findings demonstrate that overexpression labelling strategies significantly increase apparent node and paranode sizes, indicating these domains are sensitive to protein abundance, as reported for the axon initial segment (Dumitrescu et al., 2016; Galiano et al., 2012; Hedstrom et al., 2007). Our novel knock-in lines circumvent these artefacts and enable live-imaging analyses of nodes of Ranvier and paranodes closer to their native state.

### Nodes undergo morphological refinement during development

We then employed longitudinal analysis of endogenous neurofascins in our knock-in lines to monitor how axonal domains mature along the same individual peripheral nerve in vivo, which has not been possible so far with immunohistochemical fixed-tissue studies. We imaged the same individual *nfasca-mRuby3* or *nfascb-mEGFP* embryos at 3dpf, during the early stages of Schwann cell myelination, and 5dpf, when the pLL is more extensively myelinated, and assessed the morphology of individual mRuby3^+^ and mEGFP^+^ clusters (Fig. 5A,D).

**Figure 5.**
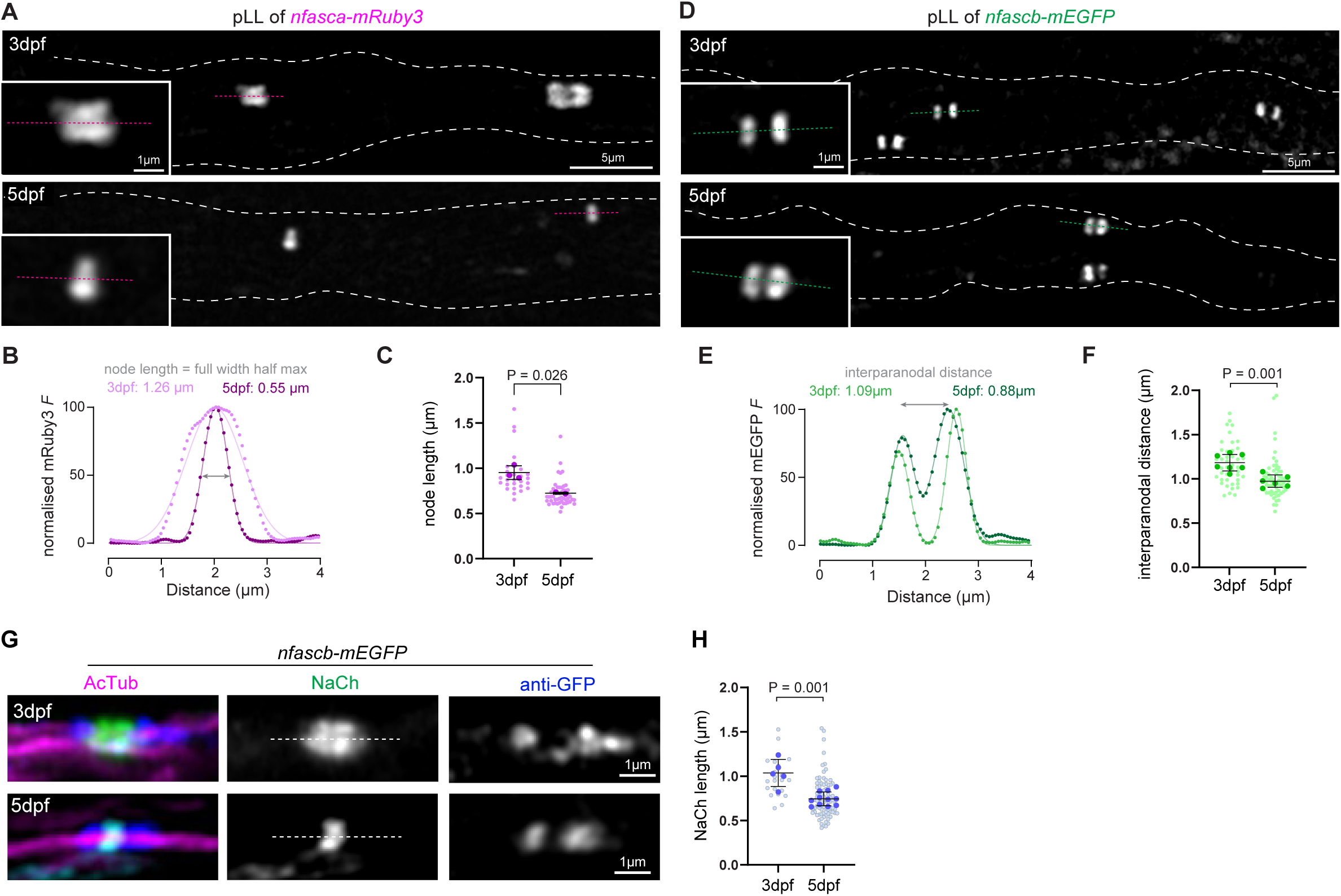
Developmental refinement of axonal domains in a zebrafish peripheral nerve. (A,D) Time-course of individual pLL nerves in *nfasca-mRuby* and *nfascb-mEGFP* at 3dpf and 5dpf. (B, E) Normalised mRuby3 and mEGFP fluorescence intensity profiles along dotted lines indicated in A and D. (C) Node length comparison between 3-5dpf: p=0.026, paired t-test, N=3 animals (F) Interparanodal distance comparison between 3-5dpf: p=0.001, paired t-test; N=7 animals (G) *nfascb-mEGFP* larvae stained for NaCh, AcTub and anti-GFP at 3dpf and 5dpf. (H) NaCh^+^ cluster length comparison: p=0.001, unpaired t-test; 5 animals at 3dpf and 11 animals at 5dpf.

In *nfasca-mRuby3* embryos, the average length of mRuby3^+^ nodal domains along pLL axons significantly decreased from 0.95±0.08µm at 3dpf to 0.72±0.01µm at 5dpf (Fig. 5B-C), a 25% reduction. In pLL axons of *nfascb-mEGFP* embryos, the length of the nodal gap between adjacent mEGFP^+^ paranodal domains significantly decreased from 1.18±0.09µm at 3dpf to 0.97±0.07µm at 5dpf (Fig. 5E-F), an 18% reduction, corroborating the shortening observed in *nfasca-mRuby3* zebrafish. We also analysed the length of individual mEGFP^+^ paranodes, which did not change significantly over time (0.56±0.04µm at 3dpf, 0.56±0.02µm at 5dpf, Fig.S2E); and compared the length of paranodes flanking the same nodal gap, which were stably asymmetric (on average 0.09µm length difference, Fig. S2F). Finally, to test if the area actually occupied by sodium channels on the axon surface also showed progressive refinement between 3-5dpf, we processed *nfascb-mEGFP* larvae for NaCh, acetylated tubulin and anti-EGFP immunohistochemistry (Fig. 5G). NaCh clusters flanked on both sides by mEGFP signal decreased in length from 1.04±0.15µm at 3dpf to 0.74±0.08µm at 5dpf, a 28% reduction (Fig. 5H).

Together, our analyses indicate that nodes of Ranvier in the pLL nerve undergo developmental shortening, similar to mammalian nerves (Panganiban et al., 2022; Smith et al., 2024), indicating that the precise morphology and position of axonal domains in peripheral nerves are refined over time.

### Schwann cells facilitate multiple steps of node development in the pLL

We next wondered whether peripheral node compaction occurred intrinsically within the axon, or involved their associated Schwann cells. First, we confirmed that Schwann cells are necessary for the initial clustering of Nfasca and Nfascb by treating larvae with the potent ErbB signaling inhibitor AG1478, an established approach to block Schwann cell migration and differentiation (Lyons et al., 2005). In *nfascb-mEGFP* larvae carrying the *Tg(nbt:DsRed)* transgene to label pLL axons treated with AG1478 from 24hpf to 4dpf, we observed a substantial reduction in mEGFP^+^ paranodal clusters in the treated nerve (Fig. 6A-B). We also treated *nfasca-mRuby3* carrying the *Tg(nbt:axon-GCaMP7s)* transgene to label pLL axons. In treated larvae, very little nodal-like mRuby3^+^ clustering was observed (Fig. 6C-D). This confirms that ErbB signaling inhibition prevents Schwann cell myelination and paranode formation, and that Schwann cells are essential for proper node assembly in zebrafish (Voas et al., 2009). This is consistent with mammalian nerves, where Schwann cells initiate clustering of NF186, concomitant with NaCh accumulation and the inward movement of flanking paranodes (Amor et al., 2017; Eshed et al., 2005; Eshed et al., 2007; Feinberg et al., 2010; Labasque et al., 2011; Malavasi et al., 2021).

**Figure 6.**
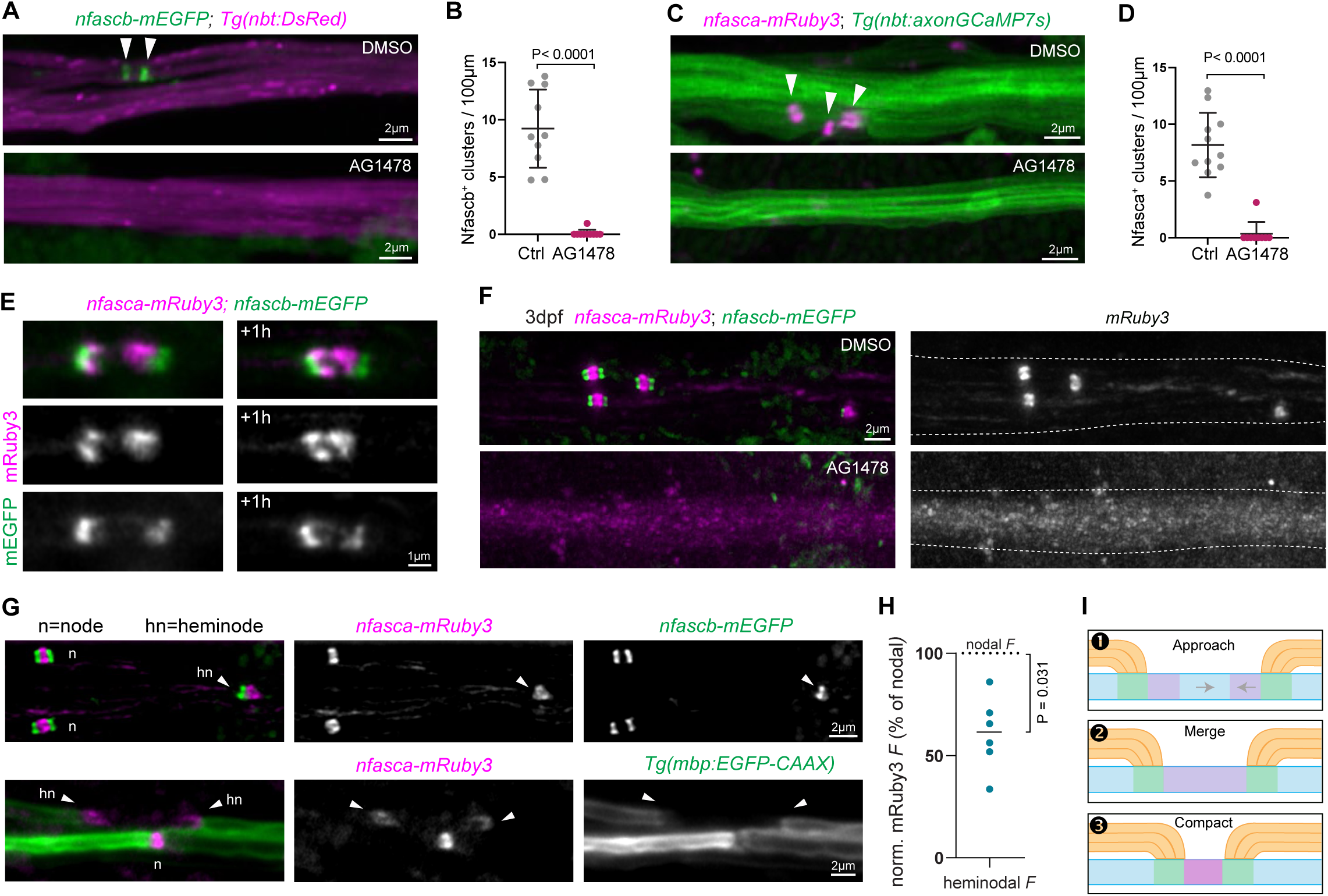
Schwann cells coordinate node and paranode assembly in the pLL. (A-B) pLL of 4dpf *nfascb-mEGFP* crossed with axonal reporter, treated with DMSO (ctrl) or AG1478. Nfascb^+^ cluster density comparison between treatments: p<0.0001, unpaired t-test with Welch’s correction, 9-11 animals. (C-D) pLL of 4dpf *nfasca-mRuby3* crossed with axonal reporter, treated with DMSO (ctrl) or AG1478. Nfasca^+^ cluster density comparison between treatments: p<0.0001, unpaired t-test with Welch’s correction, 9-11 animals. (E) Time-course of Nfasca^+^ heminodes merging into a nascent node in *(nfasca-mRuby; nfascb-EGFP)* at 3dpf. (F) pLL axons at 3dpf in *(nfasca-mRuby; nfascb-EGFP)* double knock-in treated with DMSO (ctrl) or AG1478. (G) Examples of 5dpf pLL with nodes (n) and heminodes (hn, arrowheads) present in the same field of view. (H) Heminodal mRuby3^+^ fluorescence intensity (*F*) normalised internally to the mean nodal fluorescence in the same field of view (100%= nodal *F*). P=0.031 in one-sample t-test difference from 100%, N=6 animals. (I) Model for developmental node assembly and length refinement.

We next wanted to determine how peripheral node compaction was spatiotemporally coordinated with the flanking paranodes in the same individual nerve. To simultaneously visualise nodes of Ranvier and paranodes in peripheral myelinated axons, we live-imaged double *nfasca-mRuby3; nfascb-mEGFP* larvae. Time-course analysis of the pLL in *nfasca-mRuby3; nfascb-mEGFP* larvae revealed the moment nodes of Ranvier form, when Nfascb-mEGFP^+^ paranodal clusters from opposing Schwann cells approach, concomitant with the merging of two Nfasca-mRuby3^+^ heminodes to form a typical paranode-node-paranode configuration (Fig. 6E). We never observed an mRuby3^+^ cluster that was not flanked by at least one mEGFP^+^ paranode. As expected from the single knock-in AG1478 treatments, blocking Schwann cells from developing in double *nfasca-mRuby3; nfascb-mEGFP* larvae prevented paranodes or nodes from forming, and only diffuse mRuby signal was detectable along the pLL (Fig. 6F). Since nodal gaps shorten over time (Fig. 5), and converging heminodes are flanked by tight paranodal junctions Fig. 6E), we tested if node shortening was accompanied by compaction of nodal components. We therefore assessed the fluorescence intensity of Nfasca-mRuby3^+^ clusters at heminodes vs nodes in images in which both were present, to minimise variability (Fig. 6G). In 6/6 nerves, heminodal Nfasca-mRuby3^+^ clusters (flanked by only one Nfascb-mEGFP^+^ cluster or mbp:EGFP-CAAX^+^ sheath) had lower fluorescence intensity on average compared to nodal Nfasca-mRuby3^+^ clusters in the same image (P=0.031, one-sample t-test, Fig. 6H). This suggests that after heminodes merge, nodal components compact. Together, our longitudinal imaging of converging paranodes (Fig. 5) and Schwann cell ablation experiment suggest that Schwann cells facilitate multiple steps of node development in the zebrafish pLL: clustering at heminodes, heminode convergence and merging, and nodal compaction (Fig. 6I).

### Nfascb drives post-assembly peripheral node compaction

We then wondered whether intact paranodal junctions are required to drive the nodal compaction we observed between 3-5dpf in the pLL (Fig. 5). Deleting axonal components of the paranode, such as adhesion molecules Caspr or Cntn1 or the submembranous cytoskeleton component αII-spectrin, indeed results in elongated nodes in the CNS and PNS (Amor et al., 2017; Feinberg et al., 2010; Rios et al., 2003; Rosenbluth et al., 2003; Susuki et al., 2013; Voas et al., 2007; Zhang et al., 2020), but the role of the glial paranodal component, *Nfascb/NF155*, has not been tested in the PNS. We therefore sought to abrogate Nfascb using a CRISPR-Cas9 mediated F0 knockout approach, and analyse how nodes of Ranvier developed over time in the pLL. We first tested if the nodal gap between adjacent sheaths was altered in *nfascb-mEGFP; Tg(sox10:mRFP)* larvae that were injected with *nfascb*-targeting CRISPR guides and Cas9 (Fig. 7A and *Experimental Procedures*). We readily observed sox10:mRFP^+^ sheath ends that were not mEGFP^+^ (40 of 141 sheath ends were mEGFP^+^, in 9 larvae), whereas in Cas9-only controls all sox10:mRFP^+^ sheath ends were mEGFP^+^ (117/117 sheath ends in 7 larvae). This validated the activity of our crispant strategy, disrupting the coding sequence of *nfascb* upstream of the mEGFP sequence. When we compared the length of the gap between adjacent sox10:mRFP^+^ sheath ends, we observed a significant 63% increase in mean gap length in *nfascb* crispants lacking Nfascb-mEGFP^+^ paranodes, relative to controls with both Nfascb-mEGFP^+^ paranodes at 5dpf (p=0.0096, one-way ANOVA followed by Tukey’s multiple comparisons test, Fig. 7B-D). The mosaicism of the *crispant* strategy also enabled us to ask if the presence of a single intact paranodal junction sufficed for compaction of the nodal gap. Nodal gaps in *nfascb* crispants that were flanked by either one or two Nfascb-mEGFP^+^ paranodes were not significantly longer than controls (Fig. 7D). This indicates enlarged nodal gaps in *nfascb* crispants are specifically due to paranode disruption, rather than abnormal or delayed development, and suggests even a single intact Nfascb^+^ paranodal junction provides some nodal compaction.

**Figure 7.**
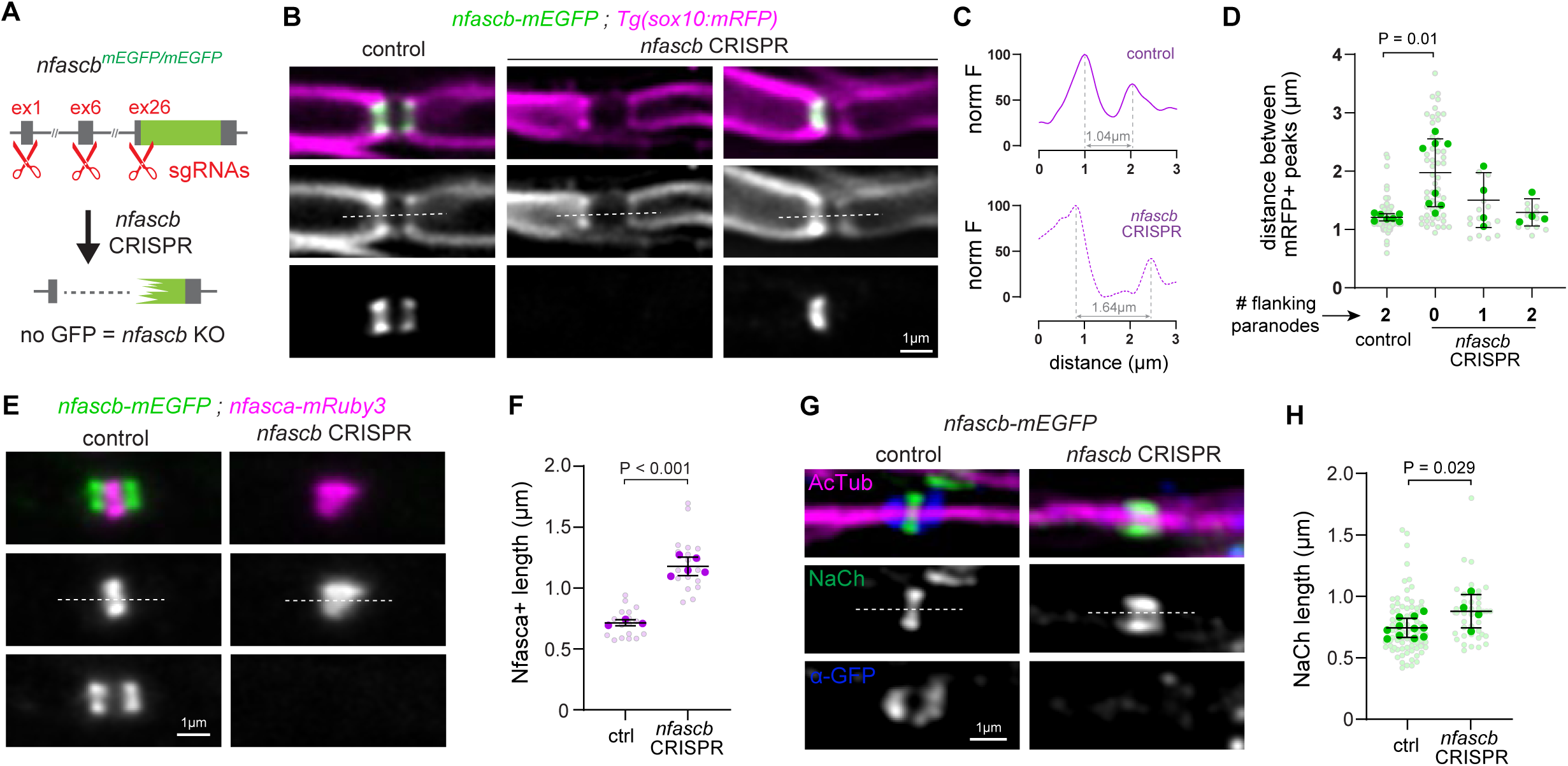
Developmental refinement of peripheral nodes requires glial Nfascb. (A) *nfascb* crispant knock-out approach targeting exons 1,6 and 26 of the *nfascb-mEGFP* locus. (B) pLL nodes in *nfascb-mEGFP; Tg(sox10:mRFP)* Cas9-only controls and crispants live-imaged at 5dpf. (C) Normalised mRFP intensity profile along nodal gap indicated in E by dotted lines. (D) pLL nodal gap comparison relative to number of paranodes present in controls and crispants: p=0.0096 for control vs no-paranode crispants, one-way ANOVA followed by Tukey’s multiple comparisons test, all other comparisons non-significant. 7 control and 8 crispant animals. (E) pLL nodes in *nfasca-mRuby3; nfascb-mEGFP* controls and crispants live-imaged at 5dpf. (F) Nfasca^+^ node length comparison: p<0.001, unpaired t-test, 3 control and 5 crispant animals. (G) pLL nodes in *nfascb-mEGFP* controls and crispants stained for NaCh, AcTub and anti-GFP at 5dpf. (H) NaCh^+^ node length comparison: p=0.028, unpaired t-test, 11 control and 4 crispant animals.

To test if the nodal domain on the axon itself was shorter, we also analysed the pLL of *nfasca-mRuby3; nfascb-mEGFP* controls and *nfascb* crispants (Fig. 7E). In controls, all 19 mRuby3^+^ node-like clusters were flanked by mEGFP^+^ paranodes (N=3 larvae), while in *nfascb* crispants, only 3 of 21 mRuby3^+^ node-like clusters were flanked by paranodes (N=4 larvae). Comparing the length of the Nfasca-mRuby3^+^ nodal clusters confirmed that, at 5dpf, nodes lacking Nfascb-mEGFP^+^ paranodes in *nfascb* crispants were on average 65% longer than control nodes (p<0.001, unpaired t-test, Fig. 7F). We then asked if the area sodium channels occupy on the axonal surface was also refined in a *nfascb*-dependent manner using immunohistochemistry (Fig. 7G). NaCh clusters in *nfascb* crispants devoid of adjacent nfascb-mEGFP signal were on average 18% longer than nodal NaCh clusters in control larvae (p=0.0285, unpaired t-test, Fig 7H). Together, our results confirm that paranodes progressively drive the compaction of all nodal components, even after the initial merging of heminodes. This identifies a non-cell-autonomous role of the glial component of paranodal junctions, Nfascb, in refining nodes of Ranvier in developing peripheral nerves.

## Discussion

Our novel nodal and paranodal knock-in reporter lines enabled the first longitudinal live-imaging analysis of endogenous neurofascins in vivo. In a developing zebrafish peripheral nerve, we revealed that after heminodes merge, nodes of Ranvier compact, and this required Schwann cells and associated paranodal adhesion, mediated by glial neurofascin Nfascb.

### How do paranodal junctions contribute to node compaction?

We showed that disrupting both paranodes flanking a node through deletion of Nfascb-mediated adhesion prevented compaction of that node. The presence of a single intact paranode provided near control levels of compaction, albeit modestly reduced; suggesting the two flanking paranodes may cooperate to compact nodes. Nodal and paranodal components such as Nfascb are tightly anchored to the underlying cytoskeleton via spectrins, ankyrins and cytoskeletal adaptor proteins (Chang et al., 2014; Ogawa et al., 2006; Zhang et al., 2013). Previous studies have shown that paranodal junctions and their associated cytoskeleton are molecular barriers restricting lateral diffusion of transmembrane proteins (Einheber et al., 2013; Horresh et al., 2010; Susuki et al., 2013; Zhang et al., 2013). Reducing the interparanodal distance, whether by the elongation of flanking myelin sheaths, local changes in the paranodal composition or in the nodal extracellular matrix, could thus induce nodal constriction increasing the density of nodal components. In addition to paranode-driven node compaction, sodium channel density may also increase through direct vesicular trafficking and insertion of channels into the nodal membrane, as proposed for the AIS (Akin et al., 2015) and peripheral nodes (Zhang et al., 2012). It will be important in future studies to determine if glial neurofascin also drives compaction in CNS nodes after their initial assembly, since NF155 and other internodal and paranodal adhesion proteins promote central myelin sheath formation and growth (Djannatian et al., 2019; Klingseisen et al., 2019; Radha et al., 2026; Zonta et al., 2008).

### Does developmental nodal compaction have functional consequences?

Using our novel reporter lines, we observed compaction in Nfasca^+^ and Nfascb^+^ reported nodal and interparanodal domains that were accompanied by refinement of voltage-gated sodium channels themselves. The observed nodal changes occur at a time when the pLL is already active and transmitting mechanosensory information to the CNS (Arancibia-Cárcamo et al., 2017; Cullen et al., 2021; Ford et al., 2015; Smith et al., 2024). Computational modelling predicts that node shortening would accelerate conduction speed, potentially by increasing sodium channel density (Klingseisen et al., 2019; Zonta et al., 2008). The morphological refinement we observed may thus represent a critical developmental mechanism to speed up conduction along this sensory nerve to refine emerging behavioural responses such as prey detection and predator evasion.

### Limitations

While our C-terminal tagging enabled analysis of neurofascins closer to their native state compared to overexpressed reporters, we cannot fully exclude that the knocked-in tags subtly disrupt protein localisation or function. This could occur through altering the precise kinetics of membrane trafficking or cytoplasmic interactions with partners. More highly-resolved analysis of single molecule behaviour, or interacting proteins, may be required to detect such defects. Additionally, in our new germ-line established knock-in lines, tagged neurofascins are expressed in all cells, which can hamper visualizing endogenous neurofascin proteins at single-cell resolution, especially in dense myelinated tracts. While less physiological than our approach, transgenic reporter overexpression enables mosaic labelling. A practical compromise for experiments that require single-cell resolution may be to carefully titrate transgene expression levels to reduce overexpression-associated artifacts by changing the number of UAS repeats (Distel et al., 2009) or using weaker cell-type specific promoters (Vagionitis et al., 2022).

### Outlook

Our live imaging observations of node and paranode dynamics raise the questions: how dynamic are individual adhesion proteins at the axon-myelin interface? What mechanisms enable such a dynamic reorganisation of these cytoskeleton-anchored domains along individual axons in vivo? Are individual adhesion complexes displaced along the axon, or are their components degraded and new complexes formed at different sites? The knock-in lines generated here provide a platform to address these outstanding gaps in our understanding of the regulation of axonal domains in vivo, in development and beyond. Recent studies in the adult nervous system show that node length can respond to changes in neuronal activity, altering nerve conduction (Arancibia-Cárcamo et al., 2017; Cullen et al., 2021), but the underlying mechanisms are not yet clear. Our finding that Nfascb controls node length during development suggests that glial regulation of paranode structure or stability could mediate changes to nodes also in adults, potentially through activity-dependent mechanisms (Osso and Hughes, 2024). Furthermore, node and paranode alterations are increasingly appreciated to occur in ageing and across a range of nervous system disorders (Cai et al., 2025; Miguel-Hidalgo et al., 2025; Ding et al., 2024; Arancibia-Carcamo and Attwell, 2014; Howell et al., 2006). Our new tools will therefore facilitate future molecular dissection of mechanisms that control the axon-myelin interface both in the healthy nervous system and in disease.

## Materials and Methods

### Zebrafish Lines and Maintenance

Zebrafish were maintained under standard conditions (Nusslein-Volhard and Dahm, 2002; Westerfield, 2000) in the BVS Aquatics facility at the Queen’s Medical Research Institute, University of Edinburgh. Studies were carried out with approval from the UK Home Office under project license PP0103366. Adult animals were kept in a 14 hours light and 10 hours dark cycle. Embryos were kept at 28.5°C in 10mM HEPES-buffered E3 embryo medium or conditioned aquarium water with methylene blue. Embryos were staged according to days postfertilisation (dpf) (Kimmel et al., 1995) and analysed up to 5dpf, before sexual differentiation. Wildtype zebrafish of the AB and WIK strains were used for knock-in experiments described in this paper. Additionally, the following established transgenic lines (Tg) were used in this study: reporters of myelination *Tg(mbp:EGFP-CAAX)* (Almeida et al., 2011) and *Tg(sox10(7.2):mRFP)* - referred to as *Tg(sox10:mRFP)* in the text (Kirby et al., 2006); pan-neuronal reporter *Tg(Xla.Tubb:DsRed)* – referred to as *Tg(nbt:DsRed)* in the text, for ‘neural-specific beta-tubulin’ (Peri and Nüsslein-Volhard, 2008); neuronal driver line *Tg(KalTA4u508)*(Antinucci et al.); and myelinating glia driver line *Tg(claudinK:Gal4)* - referred to as *Tg(cldnK:Gal4)* in the text (Münzel et al., 2012). The following knock-in lines were generated in this study: *nfasca-mRuby3*; *nfascb-mEGFP*. The following transgenic line was generated in this study: *Tg(nbt:axon-GCaMP7s)*.

### HDR template design and synthesis

Oligonucleotide primer sequences used for HDR template synthesis are indicated in Table 1. For N-terminal knock-ins for cellular expression tagging, primers included 40 nucleotide-long homology arms (italicized in Table 1) around the start codon and were designed to amplify the sequence for membrane-tethered (Fyn myristoylation motif) memEGFP or memmScarlet followed by the P2A peptide, without linker sequences. For C-terminal knock-ins for direct protein tagging, primers included 43-61 nucleotide-long homology arms around the stop codon and a GGGGS linker sequence (underlined in Table 1) in the forward primer, and were designed to amplify the sequence for mRuby3 or mEGFP, omitting the reporter start codon. The endogenous stop codon and untranslated region were used in the right homology arm. Where possible, silent alterations were introduced in the PAM sequence in the HDR template to prevent repeated cleavage (noted in bold in Table 1). Modified primers consisted of 5ʹ-end biotin (noted as ‘5Bio-’ in Table 1) and phosphorothioate bonds in the first 5 nucleotides (noted as ‘*’). All primers were obtained from IDT DNA (25-100nmole, desalted). HDR templates were produced by high-fidelity PCR from plasmids containing fluorescent protein coding sequences using Phusion or Q5 polymerase (New England Biolabs) in 50-100µL reactions. PCR products were electrophoresed on 1% agarose gels, purified using the Monarch DNA Gel Extraction Kit (New England Biolabs) and eluted in 6µL of water to maximise concentration.

**Table 1.**
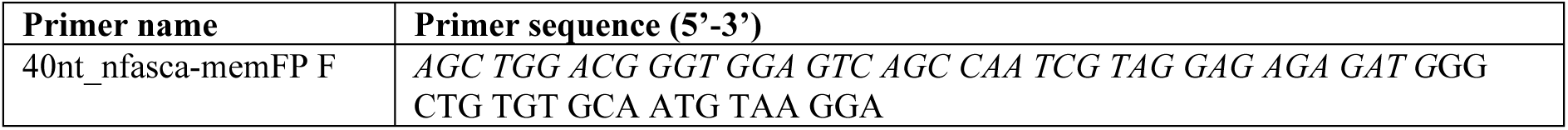
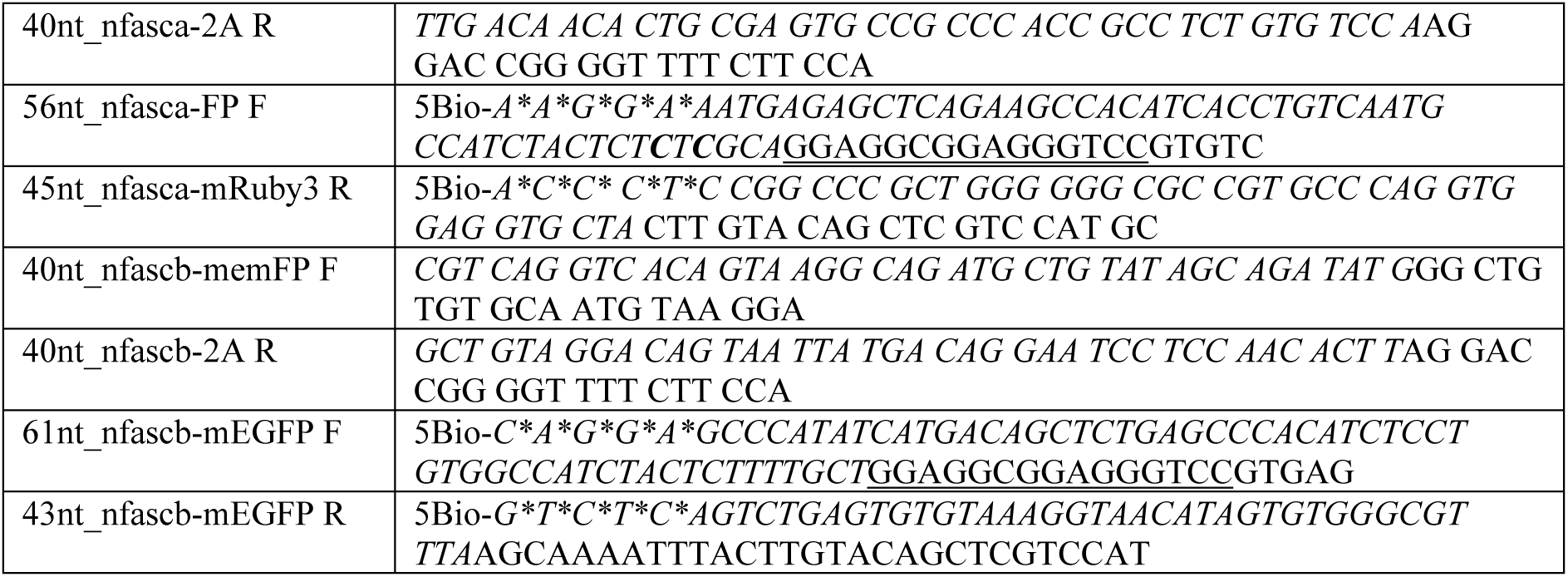
Primer sequences for HDR template synthesis.

### CRISPR reagents

CRISPR guide RNAs were designed with predicted cut sites as close as possible (≤12bp) to the desired start or stop codons (underlined in Table 2). To disrupt *nfascb* function (Fig. 7), three crRNAs targeting the start and stop codons and exon 6 were combined. crRNAs were obtained from IDT DNA (Alt-R® CRISPR-Cas9 crRNA, 2 nmol) and hybridised to universal Alt-R® CRISPR-Cas9 tracrRNA to form 20µM sgRNAs by heating to 95°C for 5min and cooling gradually to room temperature. Engen Spy Cas9 enzyme (New England Biolabs) was used.

**Table 2.**
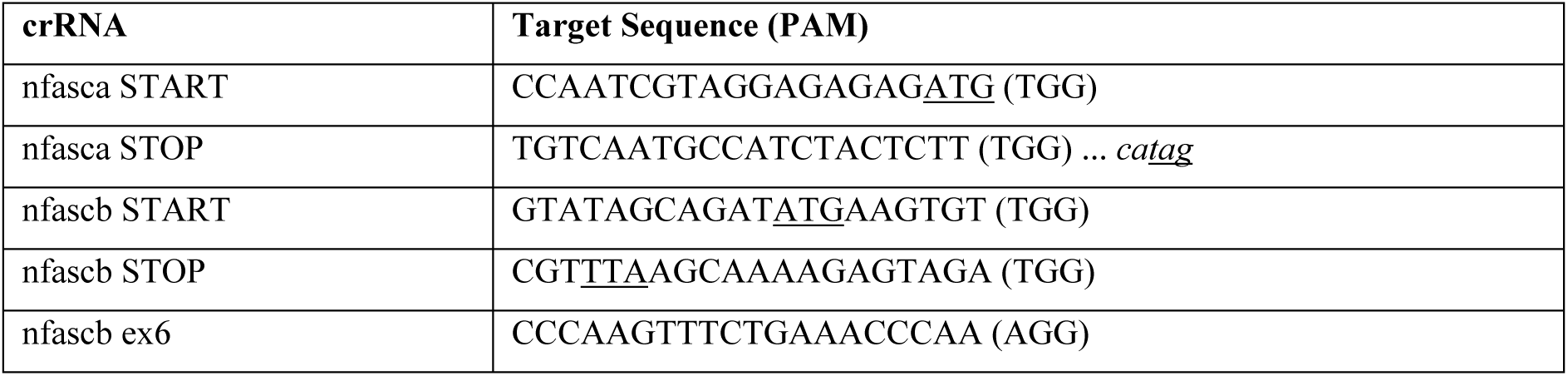
Guide RNA target sequences.

### PCR tagging solution preparation and injection

Knock-in injection solutions were prepared as before (Zhang et al., 2023) and consisted of 3-4µM sgRNA, 3-

3.2µM Engen Spy Cas9 enzyme, 10-125ng/µL of HDR template, and small-molecule HDR enhancers: 73µM NU7441 (Stratech) or 7µM HDR enhancer v2 (IDT DNA), in a final concentration of 1% DMSO. Injection solutions were prepared in a final volume of 3-10µL and incubated at 37°C for 10min to assemble the ribonucleoprotein complex (RNP), before being transferred to ice until injections took place. 1nL of solution was injected into one-cell stage wildtype (WIK or AB) eggs, directly into the cell or within the yolk as close as possible to the cell. Eggs past the one cell stage were not injected.

### Plasmid construction

The previously described expression construct pTol2-mbp:memmScarlet (Moyon et al., 2021) was used to mosaically label individual Schwann cells. To generate the Tol2 overexpression constructs pTol2-10xUAS:nfasca-mRuby3 and pTol2-10xUAS:nfascb-mEGFP, LR reactions were performed using LR Clonase II Plus (ThermoFisher Scientific) and 10fmol each of: p5E-10xUAS from the tol2kit (Kwan et al., 2007); pME-nfasca or pME-nfascb - middle-entry vectors containing the *nfasca* or *nfascb* coding sequences, p3E-mRuby3 or p3E-mEGFP; and 20fmol of destination vector pDestTol2pA2 from the tol2kit. 3-4 clones were tested for correct recombination by digestion with restriction enzymes. Subsequently, the sequences between *nfasca/b* and mRuby3/mEGFP were corrected using FastCloning (Li et al., 2011) to remove stop/start codons and *att* sequences and to introduce a short GGGGS linker, to make them identical to the knock-in alleles. The Tol2 transgenic construct pTol2-nbt:axon-GCaMP7s was recombined in an LR reaction using 10fmol each of p5E-NBT; pME-axonGCaMP7s (Almeida et al., 2021); p3E-polyA (tol2kit) and 20fmol of destination vector pDestTol2pA2 (tol2kit). 3-4 clones were tested for correct recombination by digestion with restriction enzymes. Constructs were verified by Sanger sequencing or whole plasmid sequencing (Source Bioscience). Plasmids, construction details, and sequences are available upon request.

### Microinjection

To generate transgenic lines and for mosaic labelling or overexpression in individual neurons or myelinating cells, one to two-cell stage eggs were injected with 1nL of a solution containing plasmid DNA (1-5pg/nL) and *tol2* transposase mRNA (25-50pg/nL). Fertilised embryos were sorted for normal development at the end of the injection day and then raised at 28.5°C until 2-5dpf, when they were screened for normal morphology and fluorescence.

### Knock-in germline transmission and junction analysis

F0 embryos injected with knock-in reagents and displaying fluorescence at 2-5dpf were raised to adulthood and outcrossed with wildtype strains. At least 100 F1 larvae per F0 adult were screened for fluorescence (see section ‘Microscopy’) to determine whether the F0 adult was a founder. For junction analysis, F0 or F1 animals were genotyped by first extracting genomic DNA from whole embryos between 3-5dpf using the HOTSHOT method(Meeker et al., 2007). PCR was then performed using Onetaq (New England Biolabs) with the primer pairs indicated in Table 3 and according to the manufacturer’s instructions, using 35 cycles. For this analysis, locus-specific genotyping primers were located outside of the region used for the homology arms in the HDR template. PCR products were electrophoresed in 1-2% agarose gels. Products from F0 and F1 genotyping were purified with the Monarch Gel Extraction Kit (New England Biolabs) and verified by Sanger sequencing (Source Bioscience) with the PCR primers.

**Table 3.**
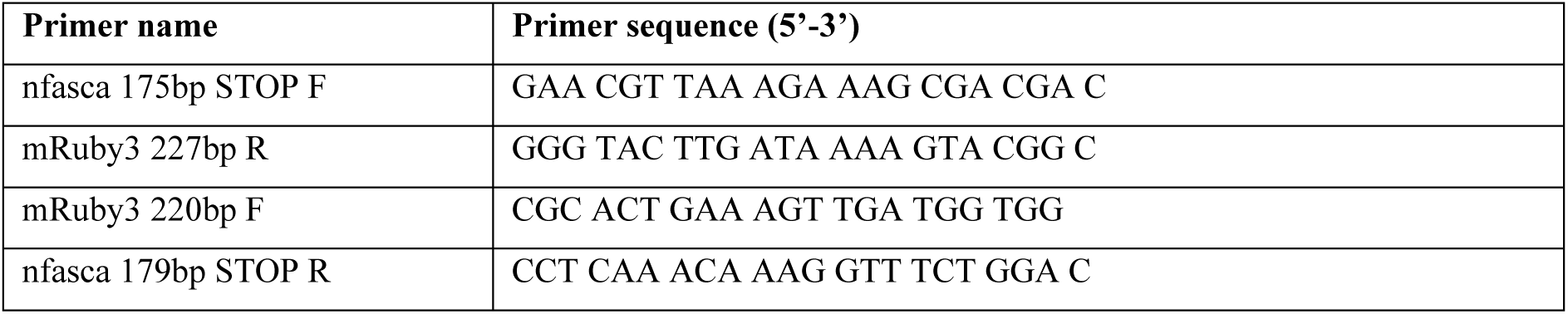
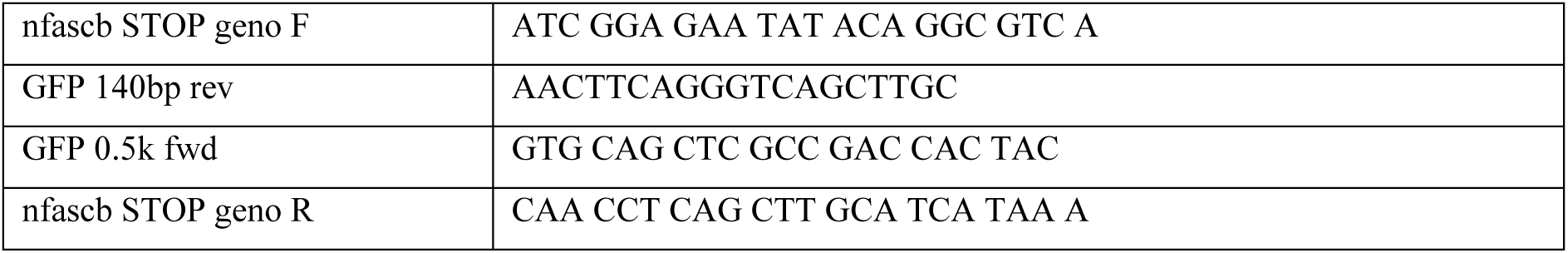
Primers for junction analysis.

### AG1478 treatment

To inhibit ErbB signaling, 2µM AG1478 in 1% dimethyl sulfoxide (DMSO) was applied to the embryo medium at 24hpf as before (Lyons et al., 2005). For efficient treatment embryos were dechorionated before application of AG1478, and embryo media with AG1478 was exchanged daily until 4dpf. As controls, 24hpf-4dpf larvae were treated with 1% DMSO applied to embryo medium.

### Immunohistochemistry

Homozygous *nfasca-mRuby3* larvae, *nfascb-mEGFP* larvae, and wild-type controls (Fig. 3); and *nfascb-mEGFP^+^ nfascb* crispant or Cas9-only control larvae obtained from outcrossing homozygous *nfascb-mEGFP* adults (Fig. 5,7) were fixed at 3-5dpf in 2% paraformaldehyde + 1% trichloroacetic acid (TCA) in phosphate buffered saline (PBS) for 12min at room temperature, followed by permeabilisation in acetone for 30min at -20°C. Samples were blocked in PBS + 1% Triton containing 10% normal goat serum for 4hrs at room temperature. Primary antibodies were diluted in 5% normal goat serum in PBS + 1% Triton and incubated overnight at 4°C. The following primary antibodies were used: mouse anti-acetylated tubulin IgG2b (Sigma-Aldrich Cat# T6793, RRID:AB_477585) at a dilution of 1:2000; mouse anti-pan Nav IgG1 (Sigma-Aldrich Cat# S8809, RRID:AB_477552) at a dilution of 1:1000; and rabbit anti-GFP (Molecular Probes Cat# A-11122, RRID:AB_221569). Secondary antibodies were diluted 1:2000 in 2% normal goat serum in PBS + 1% Triton and incubated for 4hr at room temperature, or overnight at 4°C. The following secondary antibodies were used: goat anti-mouse IgG1 conjugated with Alexa Fluor 488 (Molecular Probes Cat# A-21121, RRID:AB_2535764); goat anti-mouse IgG2b conjugated with Alexa Fluor 568 (Thermo Fisher Scientific Cat# A-21144, RRID:AB_2535780); goat anti-rabbit IgG(H+L) conjugated with Alexa Fluor 633 (Molecular Probes Cat# A-21070, RRID:AB_2535731). Samples were washed in PBS + 1% Triton and mounted in 2% low melting-point agarose on glass slides for confocal imaging as detailed below.

### Microscopy

For assessment of knock-in efficiency and germline transmission, embryos that developed with a normal morphology to 2-5dpf were first anesthetised with 600µM tricaine (3-amino benzoic acid ethyl ester, Sigma) and mounted on their sides in 1.5% low-melting point agarose in a glass coverslip, and examined individually for fluorescence along their brain and spine, using a Zeiss AxioImager A1 equipped with a Plan Apochromat 20X/0.8NA objective and a HXP120 fluorescent light source. Embryos were scored for sparse, widespread or no fluorescence. For optically-sectioned high-resolution imaging, larvae were mounted similarly and imaged at room temperature using a Zeiss LSM880 confocal with Airyscan, using 488nm (mEGFP/Alexa Fluor 488), 568nm (mRuby3/mRFP/Alexa Fluor 568) and 633nm (AlexaFluor 633) laser lines; and a Zeiss Plan-Apochromat 20X/0.8NA dry, W Plan-Apochromat 20x/1.0NA water-dipping, C-Apochromat 40x/1.2NA water-dipping or C-Apochromat 63x/1.2W Knorr UV-VIR-IR M27 water-dipping objectives. To perform photobleaching of nodes of Ranvier (Fig. S2), the ‘Regions’ function was used to define a small rectangular region-of-interest circumscribing the node, ‘Continuous’ scanning mode initiated, and the 568nm laser power briefly brought to 100% for ∼20-30 seconds until mRuby3 signal disappeared. Following acquisition, all images were Airyscan processed in 3D within ZEN Black software (Zeiss), before further processing and analysis in Fiji/ImageJ, as described below. For Fig. 2 and Supplemental Movies S1 and S2, three-dimensional renderings were generated using Arrivis Vision4D v4 software from Airyscan-processed z-stacks. Images were first converted from .czi to Arrivis .sis format. Myelin was segmented using the machine learning module, with either the mbp:EGFP-CAAX or sox10:mRFP channel used for object detection. The resulting myelin object was visualised together with the corresponding raw Nfasca-mRuby3 or Nfascb-mEGFP fluorescent signal using the 4D viewer.

### Image processing and analyses

Image analyses were performed in Fiji/ImageJ and Python aided by custom written scripts, available upon request. For comparisons of endogenous vs overexpressed neurofascins, one ∼84µm-long pLL region centered around somite 20 (*nfasca*) or two ∼105µm-long regions around somites 5-10 (*nfascb*) were imaged in each larva at 5dpf. To determine cluster length systematically while minimising inter-researcher bias, summed-intensity projections were first produced, a line ROI was first drawn along the cluster encompassing baseline background on both sides of the cluster, oriented along the underlying myelinated axon visualised by the myelination reporter. The thickness of the line was adjusted to match the width of the cluster. Then, a fluorescence intensity profile along the line was obtained, and a Gaussian curve (Nfasca-mRuby3^+^ clusters) or two Gaussian curves (Nfascb-mEGFP^+^ clusters) were fitted to the profile to remove noise. If the r^2^ measure of fit was above 0.8, the length of the cluster was determined as the full width at half maximum, calculated as 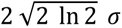. Additionally, for Nfascb-mEGFP^+^ paired clusters, the nodal gap, or inter-paranodal distance between the mEGFP^+^ peaks was determined, as the difference between the two Gaussian means. Overlapping clusters, edge clusters, or poorly fitted curves (r^2^ < 0.8) were excluded. No significant differences were observed in the length of anterior (somite 5) and posterior (somite 20) clusters, so these data were pooled, and in other experiments typically only one region was assessed.

To analyse nodes of Ranvier in the pLL after immunohistochemistry processing of homozygous knock-in larvae, at different developmental stages, and in controls vs *nfascb* crispants, one to three ∼84µm-long regions of the pLL were imaged, centered around somites 18-23. Nodes were identified as discrete NaCh^+^ clusters localised along AcTub^+^ axons and, in *nfascb-mEGFP* larvae, flanked by anti-GFP^+^ signal indicative of paranodes. To determine NaCh^+^ cluster length, local summed-intensity projections of each cluster were produced, and the full width at half maximum was calculated in a similar manner to Nfasca/b^+^ clusters.

To analyse Nfasca-mRuby3^+^ and Nfascb-mEGFP^+^ clusters in the pLL over time, two ∼66µm-long pLL regions centered around somites 5 and 20 were imaged in the same individual larvae at 3dpf and 5dpf. To compare nodal and heminodal Nfasca-mRuby3 fluorescence intensities, only animals carrying paranodal or myelination reporters were analysed to unambiguously identify nodes vs heminodes. For a strict assessment of relative intensity within-image, only images with both nodes and heminodes present were considered. Local summed-intensity projections of each cluster were produced, a line ROI was drawn along the cluster, and a fluorescence intensity profile along the line was obtained. The baseline/background was removed and a Gaussian curve was fitted. Node and heminode fluorescence intensities were determined as the amplitude of the fitted Gaussian, and averaged per nerve. Average node intensity per nerve was set at 100%, and heminode intensity expressed as % of node intensity.

To assess the formation of nodes of Ranvier and paranodes in AG1478-treated animals and controls, two ∼105µm-long pLL regions centered around somite 15 were imaged in 4dpf larvae. To assess the nodal gap between adjacent sox10:mRFP^+^ sheaths in the pLL of *nfascb* crispants and controls, two ∼84µm-long regions of the pLL were imaged, centered around somites 16-20. To determine the distance between sheaths, local summed-intensity projections of each nodal gap were produced, and the fluorescence intensity profile of a line drawn along the nodal gap was determined. The sox10:mRFP^+^ signal at the edges of myelin sheaths was not as well approximated by Gaussian curves as discrete paranodal Nfascb^+^ clusters. Therefore, the nodal gap length was determined simply as the distance between the two mRFP profile maxima which reflect sheath edges (Fig. 7B-C).

To produce figure panels, only global adjustments of brightness and contrast and background subtraction were employed. Individual z-slices, summed or maximum-intensity projections of z-stacks were used, and a representative x-y area was cropped, and transferred to Adobe Illustrator to assemble the final figures. All zebrafish images and movies represent a lateral view of the spinal cord, anterior to the left and dorsal on top, except in panels A and E in Fig. 2, which display a top-down view of the hindbrain, anterior to the top.

### Statistical Analyses

All graphs and statistical tests were produced using GraphPad Prism. Data were averaged per biological replicate for statistical comparisons (larger dots in graphs, N in figure legends indicate number of animals). Data were tested for normal distribution using D’Agostino & Pearson and Shapiro-Wilk normality tests. Normally distributed groups were compared using two-tailed unpaired or paired (Fig. 5) Student’s t-tests, or one-way ANOVA followed by Dunnet’s multiple comparisons test (Fig. 7D). Non-normally distributed groups were compared using non-parametric tests that do not assume an underlying distribution: unpaired Mann-Whitney U test, or Wilcoxon matched-pairs signed rank test as indicated in figure legends. Differences were considered significant when P < 0.05; no indication in the figures or figure legends means P > 0.05. Error bars illustrate mean ± standard deviation for normally distributed data, or median and interquartile range for non-normally distributed data.

### Data and software availability

All data supporting the findings of this study are available within the publication and its supplementary information files. All materials generated, including plasmids, knock-in and transgenic zebrafish lines will be made available upon request to the corresponding author.

## Online supplemental material

Fig. S1 shows representative Sanger sequencing chromatograms to validate precise junction integration in novel *nfasca-mRuby3* and *nfascb-mEGFP* alleles. Fig. S2 shows additional examples of Nfasca/b expression in knock-in reporter lines, including examples of time-course and time-lapse imaging. Movie S1 shows a 3D rendering of a myelinated pLL axon in *nfasca-mRuby3; Tg(mbp:EGFP-CAAX)* embryo; while Movie S2 shows a 3D rendering of myelinated pLL axon in *nfascb-mEGFP; Tg(sox10:mRFP)* embryo.

## Acknowledgements

We thank members of the Almeida laboratory, Maria Eichel-Vogel, Julia Meng, Stavros Vagionitis, and Sarah Petersen for providing feedback on the manuscript, and the Lyons and Brophy laboratories for sharing reagents. We also thank the University of Edinburgh Zebrafish Imaging and Screening Facility for microscopy support, and the University of Edinburgh BVS Aquatics Facility for zebrafish husbandry and support. This work was supported by a Chancellor’s Fellowship to R.A. from the University of Edinburgh; a Biotechnology and Biological Sciences Research Council grant (BB/X009394/1) and UK Research and Innovation grant under the UK government’s Horizon Europe Funding Guarantee (EP/Y024311/1) to R.A. S.S is a recipient of a Walter Benjamin Fellowship from the Deutsche Forschungsgemeinschaft (DFG Grant 565898524).

## Author Contributions

Conceptualisation, P.L-B, K.M-P, S.S and R.A. Investigation, P.L-B, K.M-P, S.S, M.P, and R.A. Writing - Original Draft, P.L-B, K.M-P, S.S and R.A. Writing – Review & Editing, P.L-B, K.M-P, S.S, M.P and R.A. Supervision, Project Administration, and Funding Acquisition, R.A.

---

**Figure S1.**
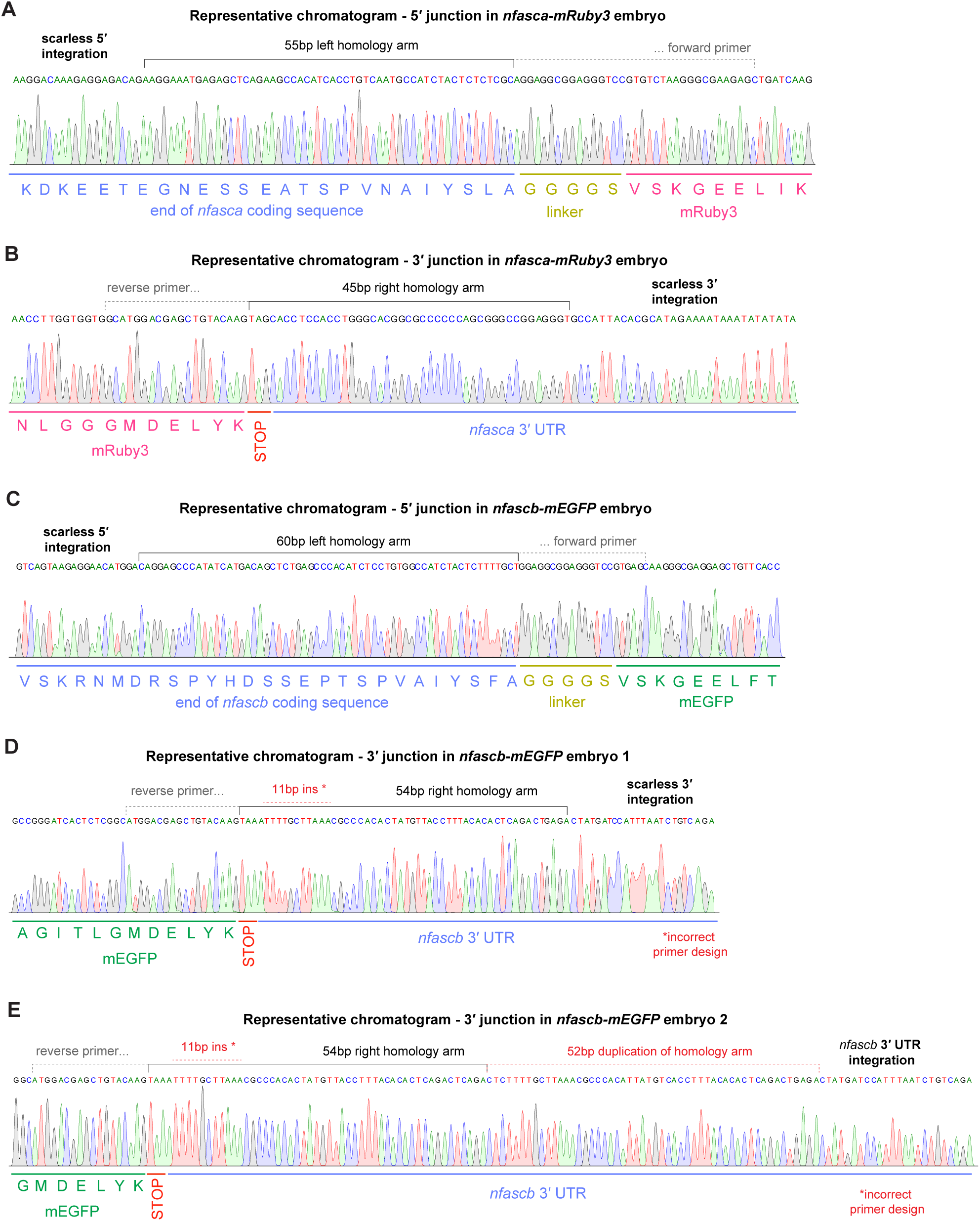
Sanger sequencing validation of novel *nfasca-mRuby3* and *nfascb-mEGFP* alleles. (A-B, C-E) Representative chromatograms from 5’ and 3’ junction Sanger sequencing of alleles in individual *nfasca-mRuby3* and *nfascb-mEGFP* F1 or F2 embryos. Note that the 3’ junction of *nfascb-mEGFP* embryos contains a 11bp insertion after the stop codon compared to the wildtype genomic sequence due to incorrect primer design; this does not affect the protein sequence. We also recovered a second allele with a partial duplication of the homology arm in the 3’ UTR (E).

**Figure S2.**
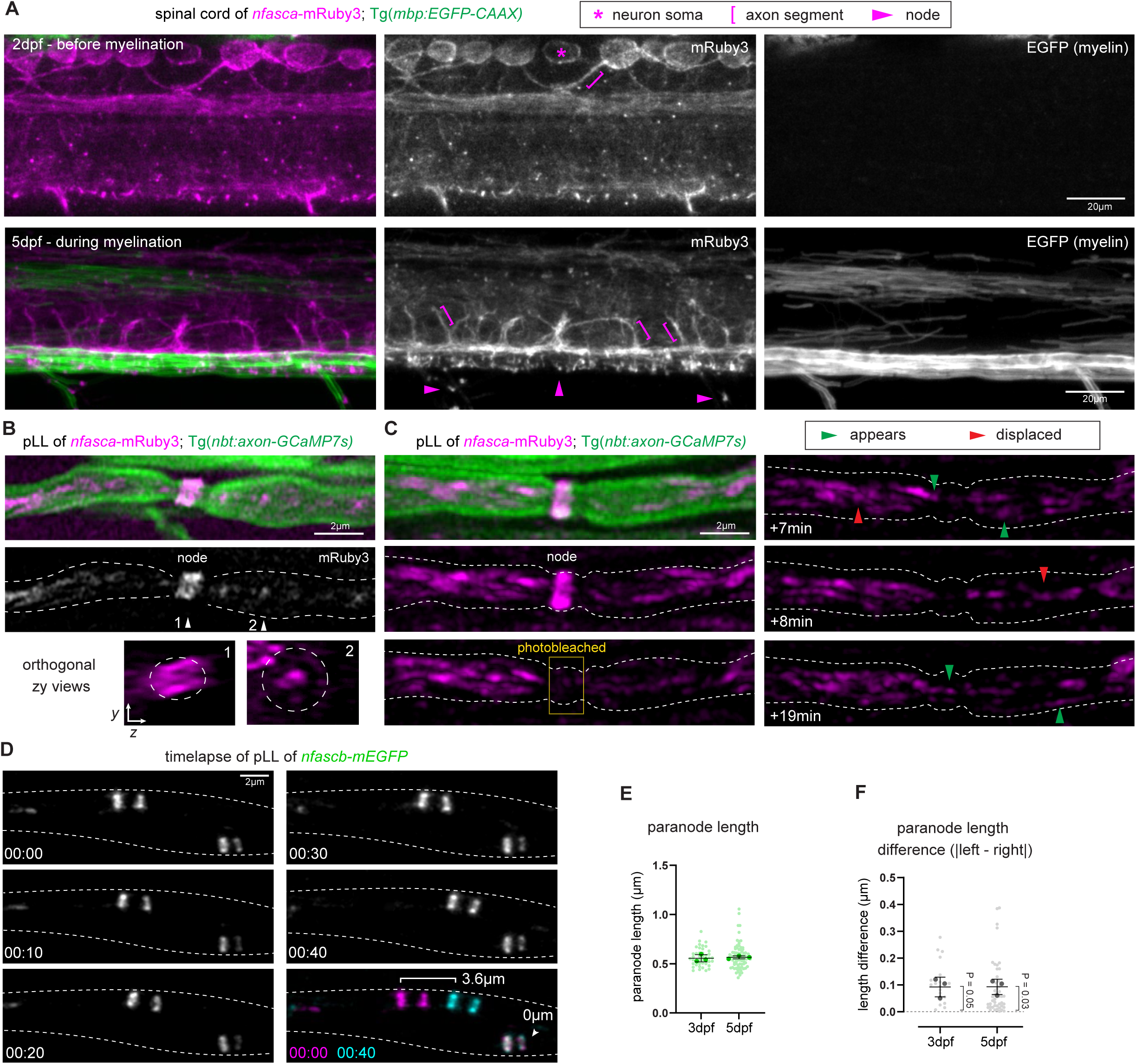
Novel *nfasca/b* knock-in lines enable analyses of axonal domain dynamics. (A) Live-imaging of the spinal cord before (2dpf) and after (5dpf) the onset of myelination. Asterisk indicates Nfasca signal present in Rohon-Beard neuron-like somas (e.g. asterisk) and unmyelinated axonal segments (e.g. bracket). At 5dpf, Nfasca signal is present in putative axon initial segments (e.g. brackets) and nodes of Ranvier (e.g. arrowheads). (B-C) High-magnification live-imaging of pLL axons. Orthogonal zy views and photobleaching help discern nodal (1) and non-nodal (2) Nfasca and its trafficking. Green arrowheads indicate examples of Nfasca^+^ particles that appeared since the previous frame, and red arrowheads indicate Nfasca^+^ particles that are displaced in the next frame. (D) Timelapse imaging showing node-paranode assemblies can be rapidly displaced along pLL axons. (E) pLL paranode length comparison between 3-5dpf (p=0.623, paired t-test). (F) Quantification of the left-right paranode length difference in a nodal assembly between 3-5dpf (p=0.049 at 3dpf and p=0.030 at 5dpf, one-sample t-test of difference from zero).

**Supplementary Movie S1** – 3D rendering of myelinated pLL axon in *nfasca-mRuby3; Tg(mbp:EGFP-CAAX)* embryo. Myelin in green, Nfasca-mRuby3 signal in magenta.

**Supplementary Movie S2** – 3D rendering of myelinated pLL axon in *nfascb-mEGFP; Tg(sox10:mRFP)* embryo. Myelin in magenta, Nfascb-mEGFP signal in green

